## Supplementary figures and images for "Systematic identification of intron retention associated variants from massive publicly available transcriptome sequencing data"

### Supplementary Figure 1

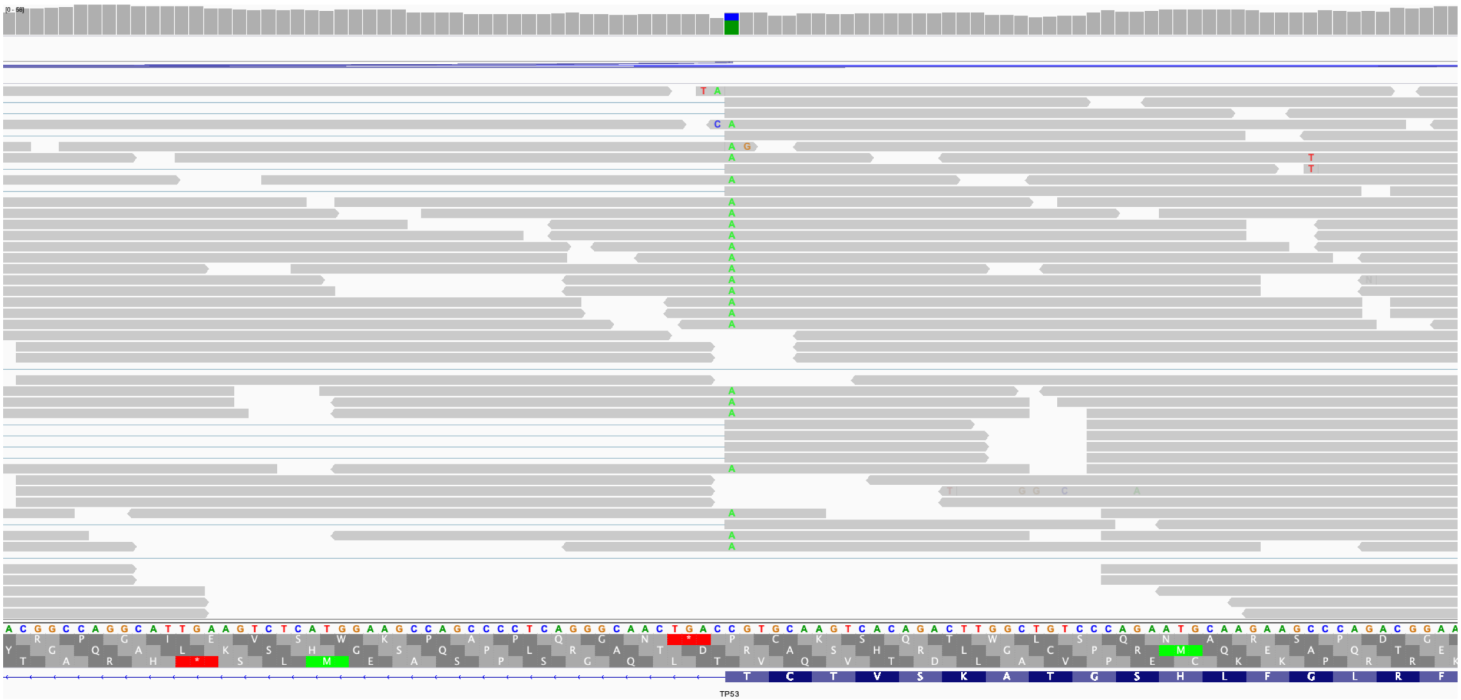

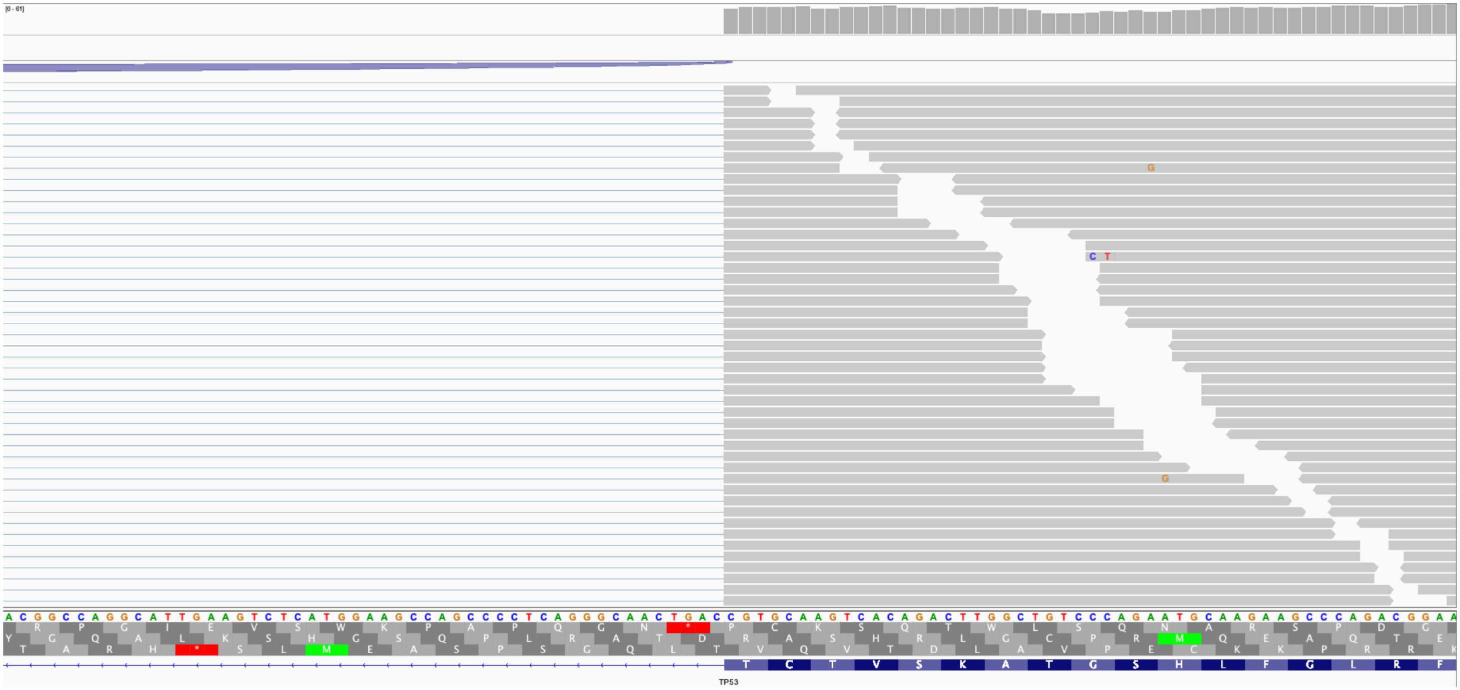

### Supplementary Figure 2

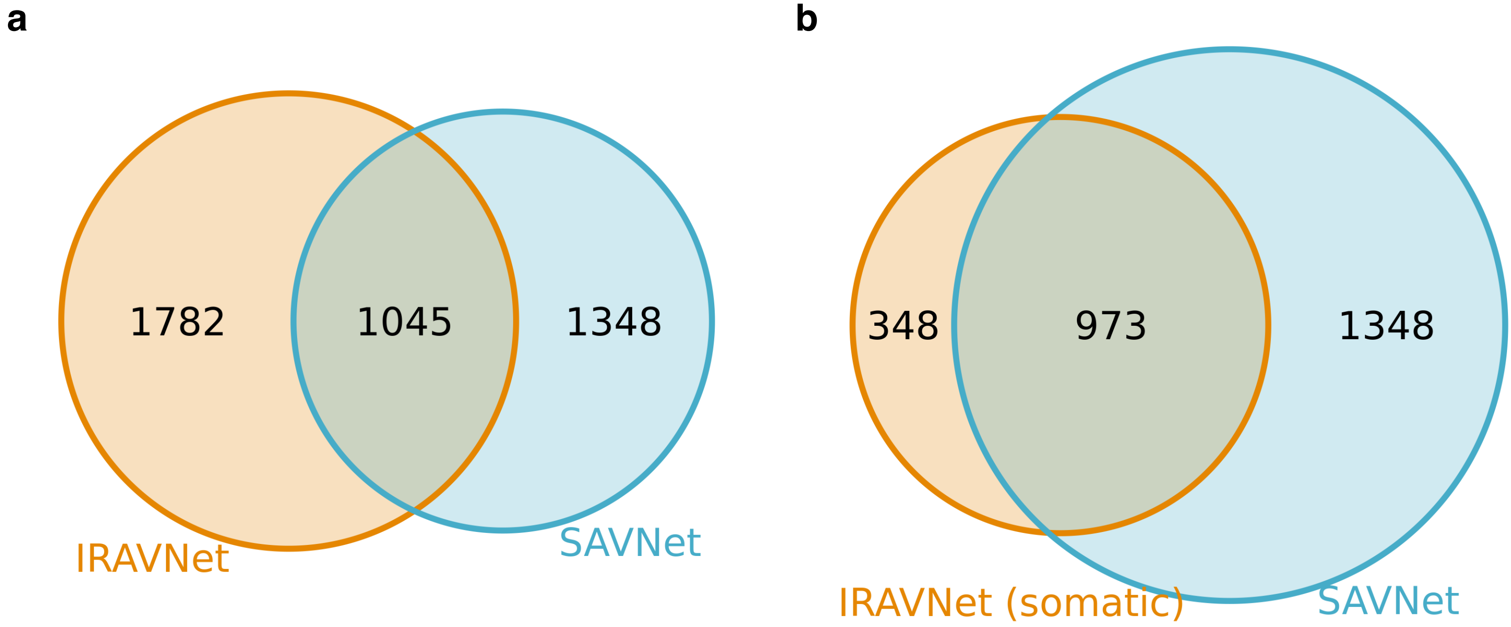

### Supplementary Figure 3

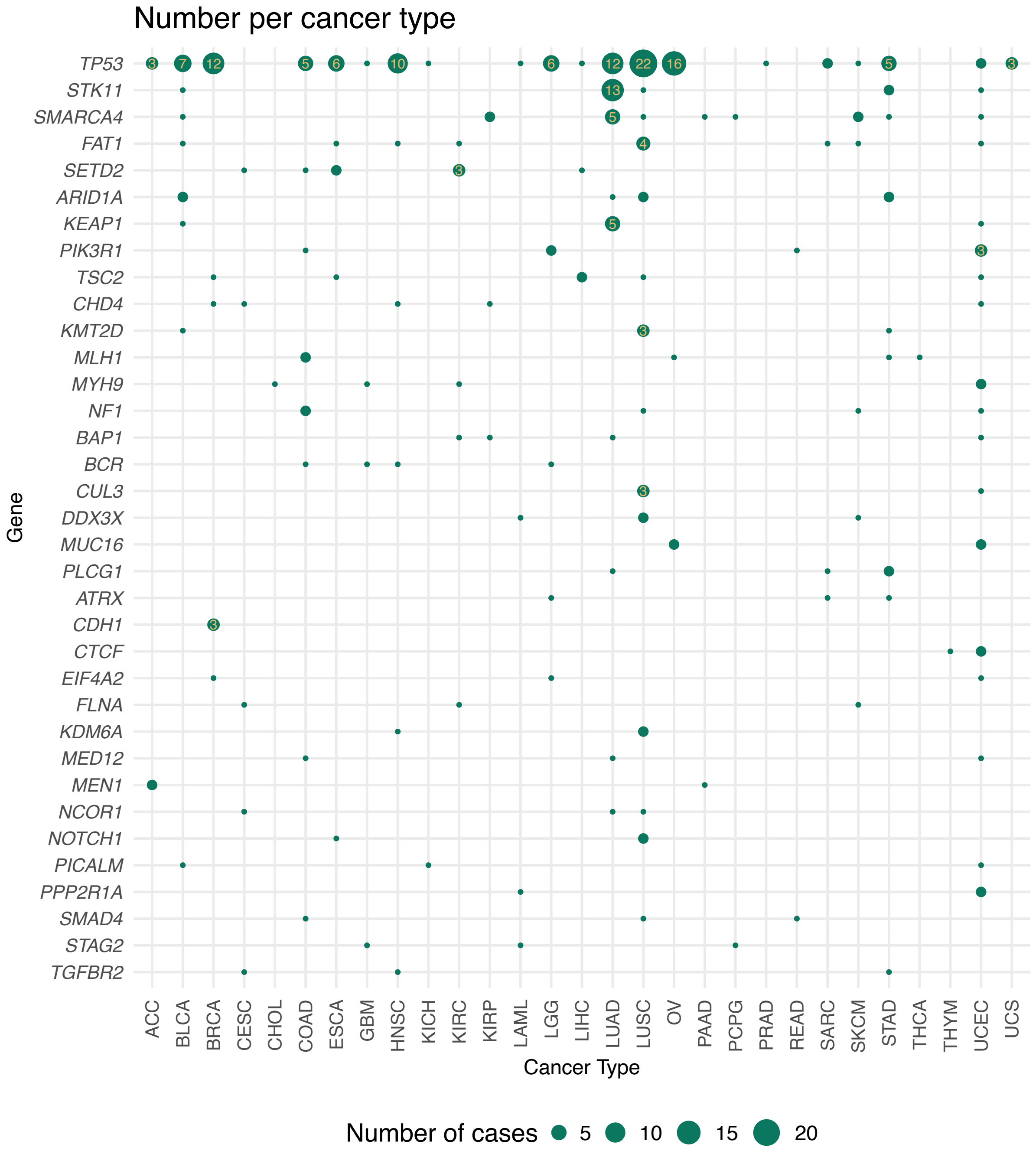

### Supplementary Figure 4

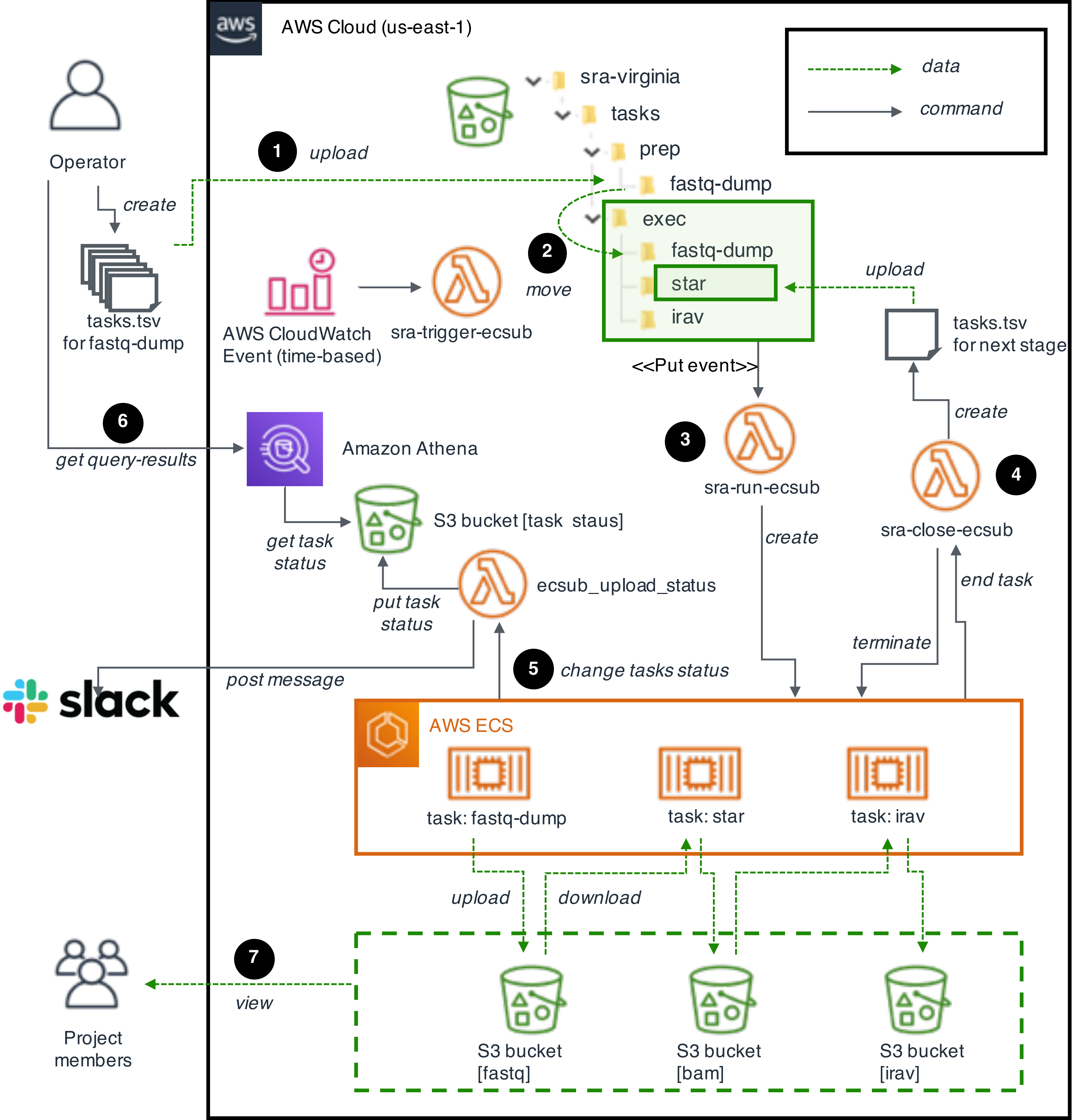

### Supplementary Figure 5

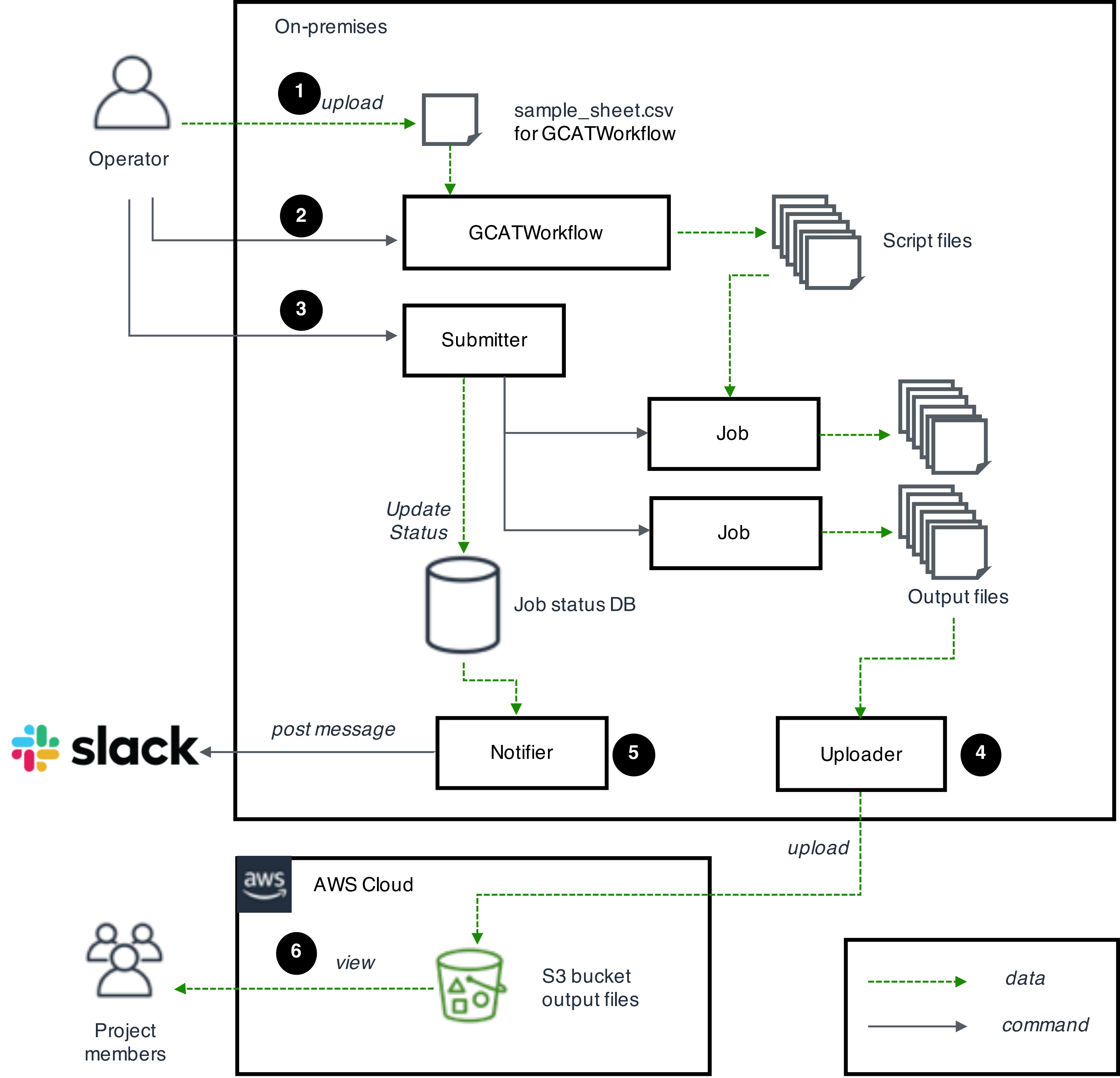

### Supplementary Figure 6

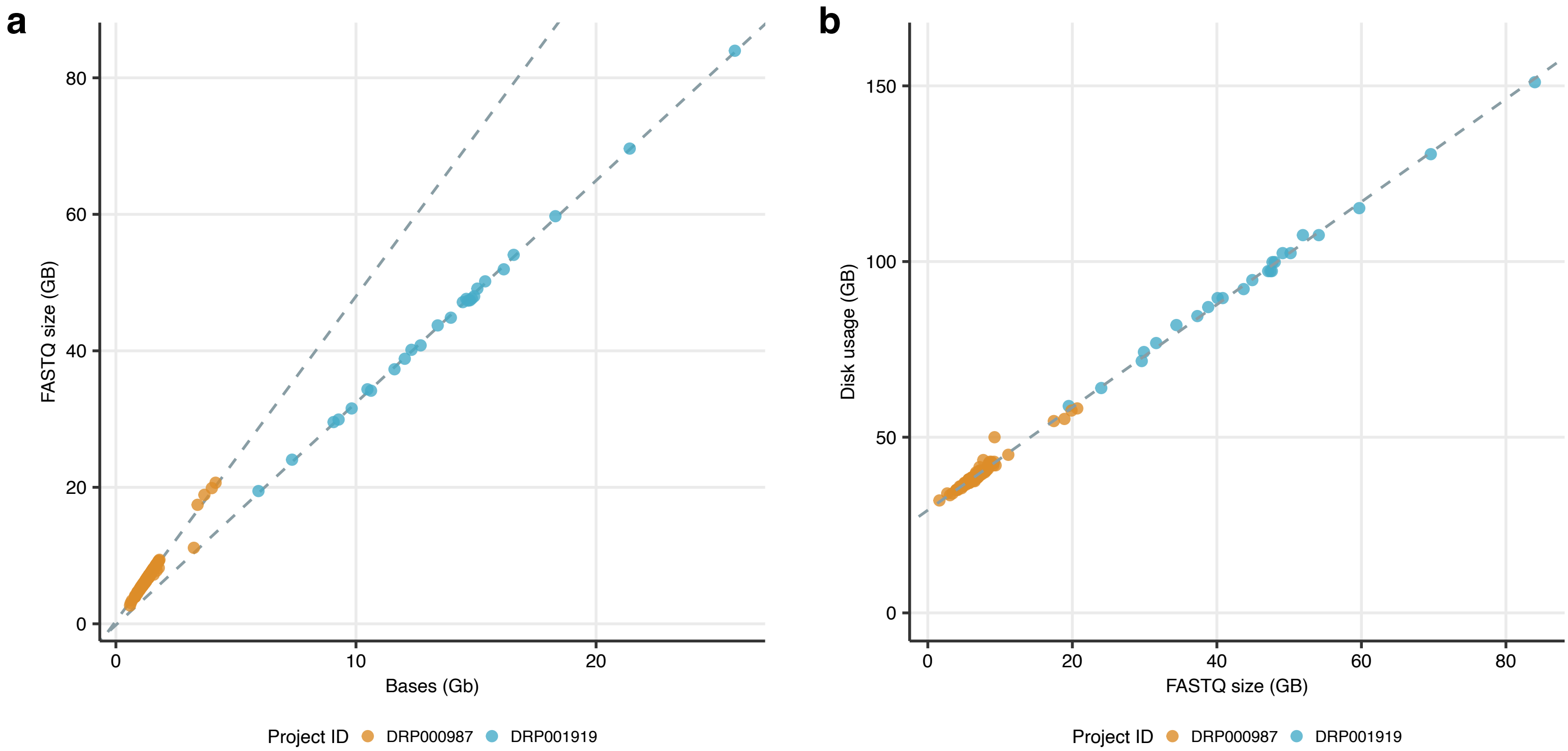

### Supplementary Figure 7

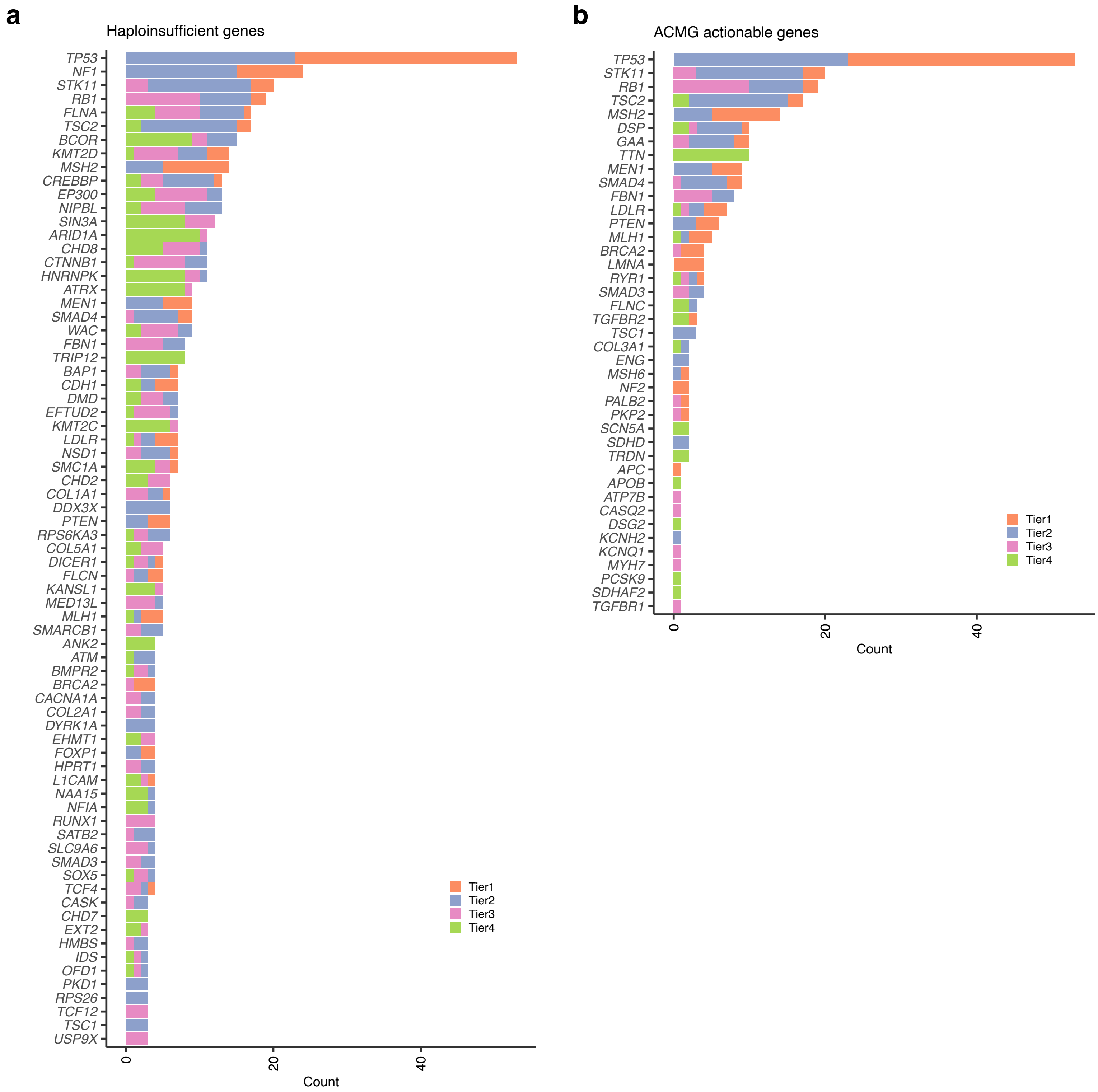

### Supplementary Figure 8

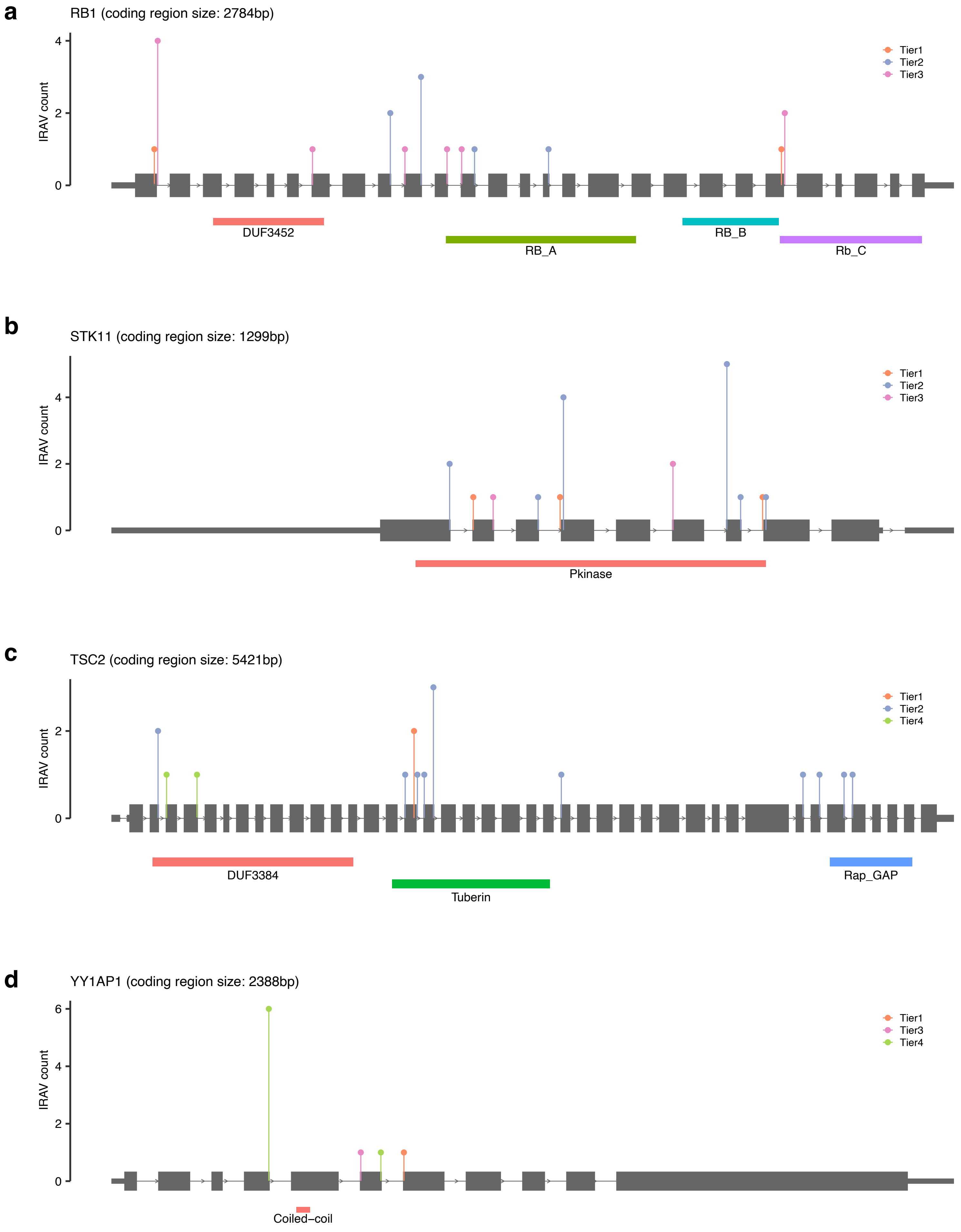

### Supplementary Figure 9

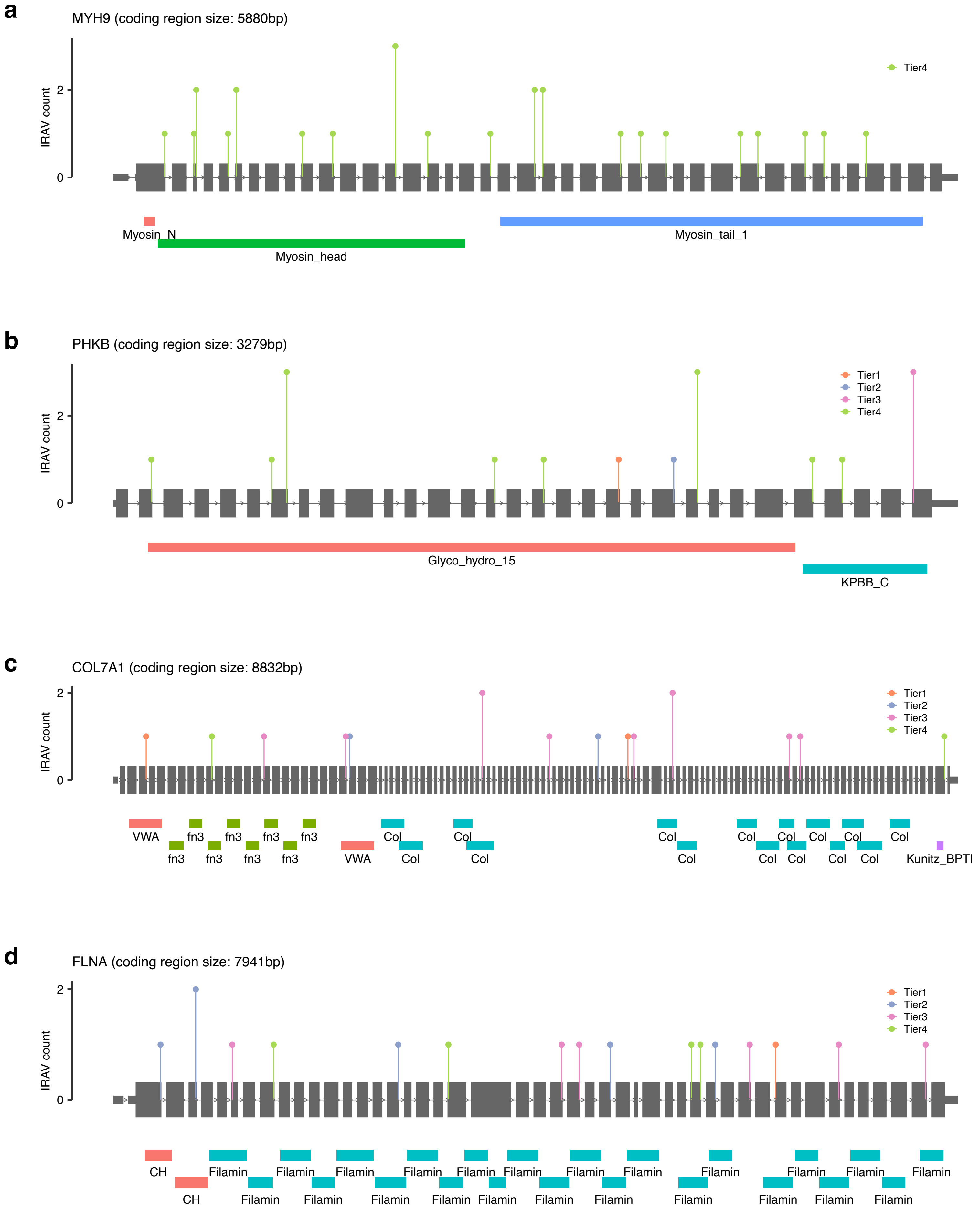

### Supplementary Figure 10

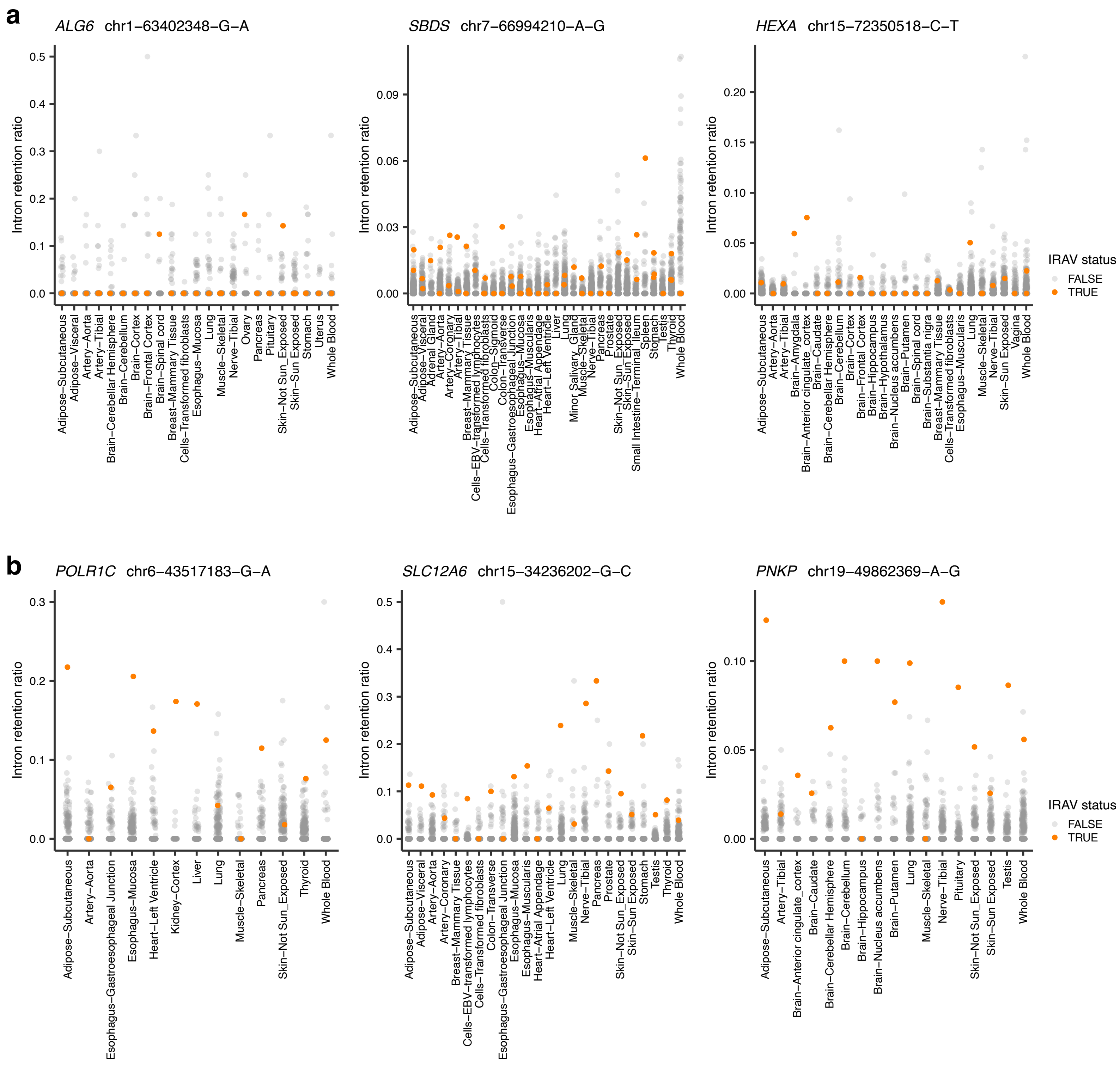

### Supplementary Figure 11

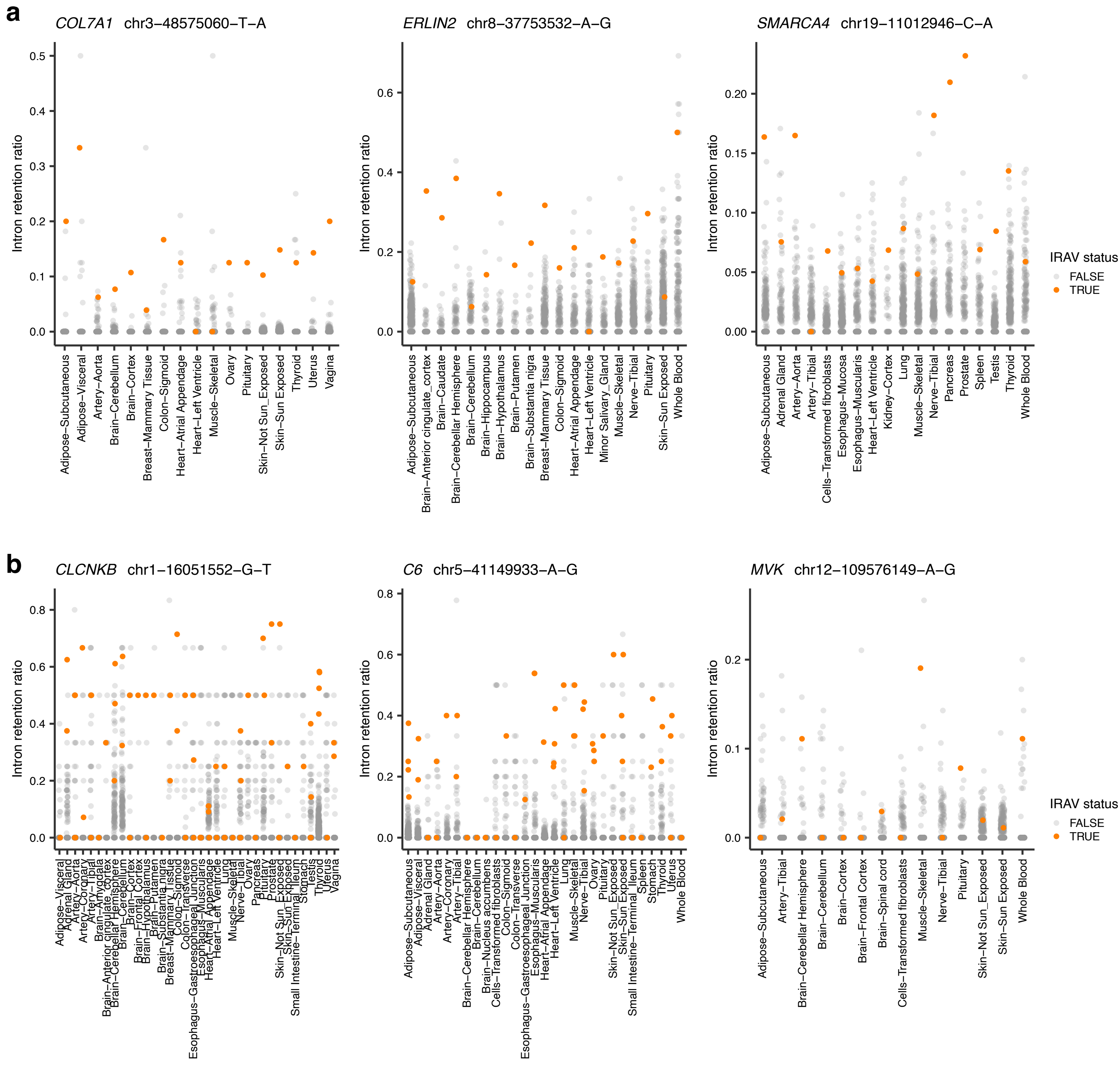
